# Drug combinations with apoptosis pathway targeted agents alrizomadlin, pelcitoclax, and dasminapant in multi-cell type tumor spheroids

**DOI:** 10.1101/2024.06.11.598557

**Authors:** Nathan P. Coussens, Thomas S. Dexheimer, Thomas Silvers, Phillip R. Sanchez, Naoko Takebe, James H. Doroshow, Beverly A. Teicher

## Abstract

Apoptosis, or programmed cell death, plays a critical role in maintaining tissue homeostasis by eliminating damaged or abnormal cells. Dysregulation of apoptosis pathways is a hallmark of cancer, allowing malignant cells to evade cell death and proliferate uncontrollably. Targeting apoptosis pathways has emerged as a promising therapeutic strategy in cancer treatment, aiming to restore the balance between cell survival and death. In this context, the MDM2 inhibitor alrizomadlin, the Bcl-2/Bcl-xL inhibitor pelcitoclax, and the IAP family inhibitor dasminapant were evaluated both individually and in combination with standard of care and investigational anticancer small molecules with a spheroid model of solid tumors. The multi-cell type tumor spheroids were grown from endothelial cells and mesenchymal stem cells combined with human malignant cells that were either established or patient-derived cell lines from the NCI Patient-Derived Models Repository. The malignant cell lines were derived from a range of solid tumors including uterine carcinosarcoma, synovial sarcoma, rhabdomyosarcoma, soft tissue sarcoma, malignant fibrous histiocytoma, malignant peripheral nerve sheath tumor (MPNST), pancreas, ovary, colon, breast, and small cell lung cancer. Interactions were observed from combinations of the apoptosis pathway targeted agents. Additionally, interactions were observed from combinations of the apoptosis pathway targeted agents with other agents, including PARP inhibitors, the XPO1 inhibitor eltanexor, and the PI3K inhibitor copanlisib. Enhanced activity was also observed from combinations of the apoptosis pathway targeted agents with MAPK pathway targeted agents, including the MEK inhibitor cobimetinib as well as adagrasib and MRTX1133, which specifically target the KRAS G12C and G12D variants, respectively.

**SIGNIFICANCE:** Multi-cell type tumor spheroids grown from normal and patient-derived malignant cell lines were screened to identify potentially efficacious combinations with the investigational agents alrizomadlin, pelcitoclax or dasminapant that target cell death pathways. This article highlights novel combinations with FDA approved drugs including eltanexor, cobimetinib and copanlisib.

## Introduction

Cell death is a highly regulated normal process in the development and maintenance of complex organisms. Among the numerous mechanisms of cell death, apoptosis is foremost in maintaining a balance between cell death and division. Failure to undergo apoptosis, or evading its regulatory mechanisms, leads to aberrant cellular proliferation, a hallmark of cancer (1). In addition, defects in apoptosis pathways can contribute to chemotherapy resistance (2). Thus, there has been significant interest in the development of apoptosis targeted anticancer drugs (3).

In cancer cells, the disruption of apoptosis regulators within both the extrinsic and intrinsic apoptotic pathways commonly includes the tumor suppressor protein p53, anti-apoptotic B-cell lymphoma 2 (Bcl-2) proteins, and inhibitors of apoptosis proteins (IAPs), making them viable drug targets (4). The tumor suppressor gene TP53 that encodes the p53 protein is the most frequently altered gene in human tumors (∼50%) (5). The p53 protein acts as a transcription factor, initiating transcription for proteins involved in apoptosis, senescence, and repair in response to various cellular stresses, including DNA damage, hypoxia, and nutrient deprivation (6). The loss of p53 function due to mutation or deletion can prevent entry of damaged cells into apoptosis, consequently leading to the development of malignancy (7,8). In normal cells, a low level of the p53 protein is maintained by the E3 ubiquitin ligase MDM2 (murine double minute 2), which targets p53 for proteasomal degradation to facilitate its continuous elimination. Because MDM2 functions as a negative regulator of p53, inhibiting their interaction presents a potential therapeutic strategy for restoring p53 function (9,10). Alrizomadlin (APG-115) is a recently developed inhibitor that targets the p53/MDM2 interaction, thereby restoring the tumor suppressor function of p53 and initiating apoptosis in tumor cells harboring wildtype p53 (11).

Another central regulator of the intrinsic apoptotic pathway is the Bcl-2 family of proteins. Within this family, there are members that promote cell survival, such as Bcl-2 and Bcl-xL, and others that induce apoptosis, such as BIM, BAD, BAX, and BAK. The proapoptotic proteins both interact with and sequester the antiapoptotic Bcl-2 proteins, while also activating downstream effectors. This cascade leads to mitochondrial permeabilization, release of cytochrome-c, and activation of caspases, which initiates apoptosis (12). Maintaining the balance of Bcl-2 proteins is critical for apoptosis induction, thus strategies to modulate their expression or interactions hold potential anticancer utility (13–16). The first generation of Bcl-2 family inhibitors, designed as BH3 mimetics, included the highly selective Bcl-2 inhibitor ABT-199 (venetoclax), which was approved by the FDA for treatment of hematological malignancies, either as a standalone therapy or in combination. Venetoclax works by displacing proapoptotic BH3-only proteins from Bcl-2 and triggering the apoptosis cascade (17,18). Pelcitoclax (APG-1252) is a second-generation BH3 mimetic designed to selectively inhibit Bcl-2 and Bcl-xL. This compound was formulated as a prodrug to circumvent the on-target toxicity observed in platelets with previous agents that targeted Bcl-xL (19–21).

Lastly, IAPs serve as a family of negative regulators that influence caspases and cell death mechanisms through both intrinsic and extrinsic pathways. Elevated levels of IAP proteins contribute to enhanced cell survival, chemo-resistance, disease progression, metastasis, and poor prognosis (22,23). Endogenous antagonists of IAPs, such as the second mitochondria-derived activator of caspases (Smac/DIABLO), interact with the BIR domains of IAPs, disrupting their ability to bind to and inhibit caspases (24–26). Birinapant, a first-generation Smac mimetic IAP inhibitor, interacts with two IAP family members, XIAP and cIAP1, inducing cell death and triggering cIAP1 degradation and caspase activation (27,28). Dasminapant (APG-1387) is a more advanced bivalent Smac mimetic that exhibited anti-tumor effects in various cancer cell lines and xenograft tumor models (29–32) and is presently undergoing clinical trials.

This study evaluated three investigational agents that modulate cell death pathways and were designed to restore apoptosis by inhibiting diverse targets. The MDM2 inhibitor alrizomadlin, the Bcl-2/Bcl-xL inhibitor pelcitoclax, and the IAP inhibitor dasminapant were utilized to explore the activity of apoptosis pathway targeted agents in combination with other targeted agents. The activities of single agents and combinations were evaluated in a multi-cell type tumor spheroid model developed from patient-derived cell lines of the National Cancer Institute (NCI) Patient-Derived Models Repository (PDMR, https://pdmr.cancer.gov/) and several well-established human malignant cell lines. The malignant cell lines spanned a range of cancer types, including uterine carcinosarcoma, synovial sarcoma, rhabdomyosarcoma, soft tissue sarcoma, malignant fibrous histiocytoma, malignant peripheral nerve sheath tumor (MPNST), pancreas, ovary, colon, breast, and small cell lung cancer (SCLC).

## Materials and Methods

### Compounds

The drugs and investigational agents: alrizomadlin (APG-115; NSC831270), pelcitoclax (APG-1252; NSC835773), dasminapant (APG-1387; NSC834197), TAK-243 (MLN7243, NSC785004), copanlisib (NSC816437), olaparib (NSC753686), talazoparib (NSC767125), cobimetinib (NSC768068), eltanexor (KPT-8602, NSC794443), abemaciclib (NSC768073), sotorasib (NSC818433), adagrasib (NSC831453), MRTX1133 (NSC836407), Iadademstat (ORY-1001, NSC806812), and RMC-4630 (NSC839698), were obtained from the NCI Developmental Therapeutics Program Chemical Repository. The FDA-approved anticancer drug set is available from the Developmental Therapeutics Program at https://dtp.cancer.gov/organization/dscb/obtaining/. The drugs and investigational agents used in this study were demonstrated to be >95% pure by proton nuclear magnetic resonance and liquid chromatography/mass spectrometry. The stock solutions were prepared in dimethyl sulfoxide (DMSO, Sigma-Aldrich, St. Louis, MO, cat. D2650) at 800-fold the tested concentration and stored at −70 °C prior to their use. All drugs and investigational agents were tested over a range starting from a high concentration at or near the clinical C_max_ and decreasing in half-log increments. If the clinical C_max_ for an agent had not been determined, the highest concentration tested was 10 µM (**Table 1**).

**Table 1.**
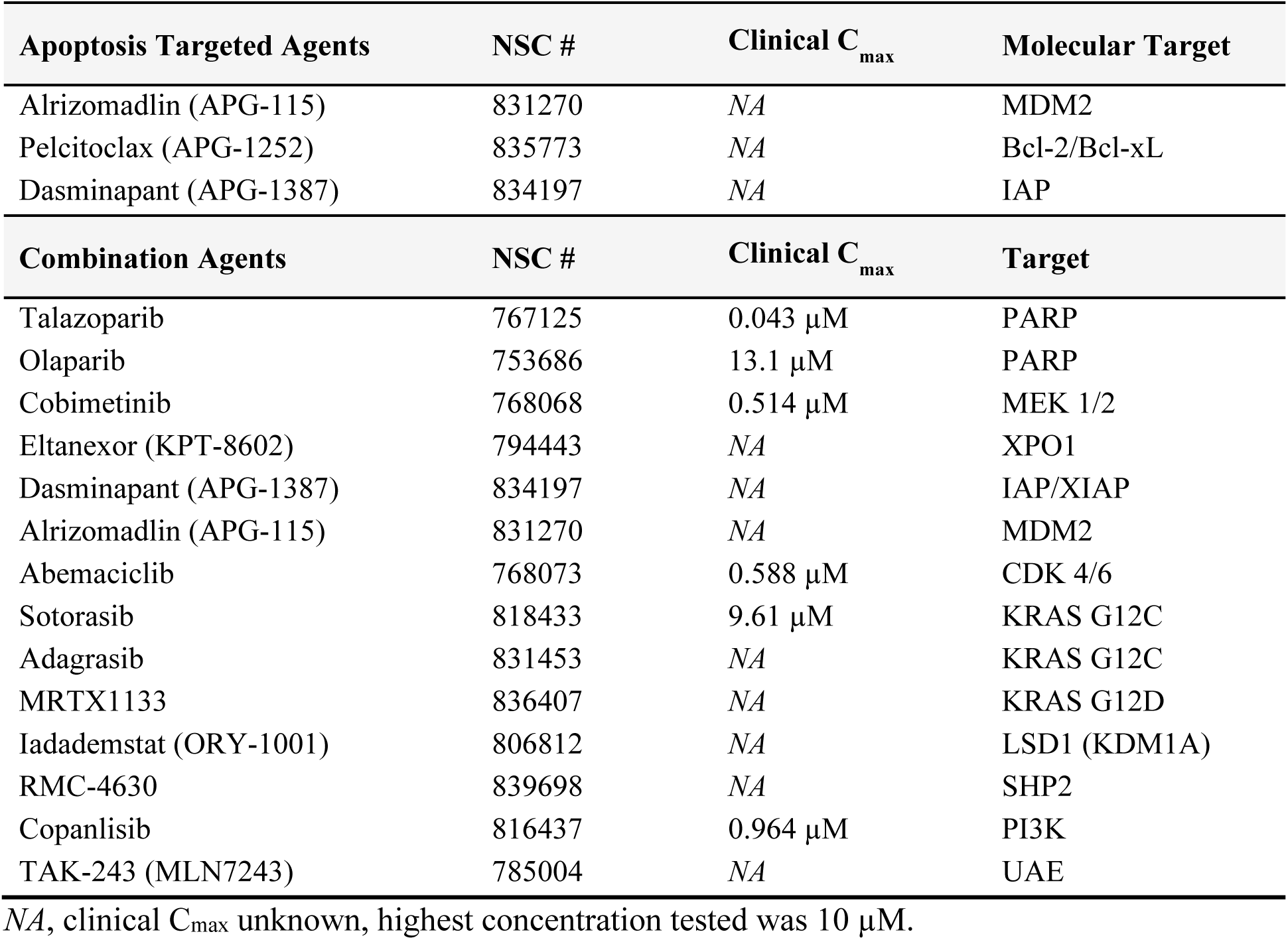
FDA-approved drugs and investigational agents used in this study. The study included apoptosis targeted agents (*top*) and molecular targeted combination agents (*bottom*). If available at the time of this study, the clinical C_max_ is listed.

### Cell Lines

The patient-derived cancer (PDC) cell lines were obtained from the NCI PDMR (https://pdmr.cancer.gov) and included five colon adenocarcinomas: 186277-243-T-J2-PDC, 439559-082-T-J2-PDC, 463943-066-T-J2-PDC, 519858-162-T-J1-PDC, and 616215-338-R-J1-PDC; three pancreatic ductal adenocarcinomas: 323965-272-R-J2-PDC, 521955-158-R2-J5-PDC and 521955-158-R6-J3-PDC; two uterine carcinosarcoma: 327498-153-R-J2-PDC and 993429-296-R-J1-PDC; two rhabdomyosarcomas: 755483-174-R-J1-PDC and 772245-204-R-J1-PDC; two soft tissue sarcomas: 126254-015-R-J1-PDC and 494315-158-R-J1-PDC; two malignant fibrous histiocytomas: 317291-083-R-J1-PDC and 596521-263-R-J1-PDC; two invasive breast carcinomas: 171881-019-R-J1-PDC and 885512-296-R-J2-PDC; and one ovarian clear cell carcinoma: 556581-035-R-J1-PDC. The established synovial sarcoma cell lines HS-SY-2 (RRID:CVCL_8719) and SYO-1 (RRID:CVCL_7146), as well as the malignant peripheral nerve sheath tumor cell line MPNST were obtained from the Sloan Kettering Institute. The following established cell lines were purchased from the American Type Culture Collection (ATCC, Manassas, VA): PANC-1 pancreatic carcinoma (ATCC cat. CRL-1469, RRID:CVCL_0480), MCF7 invasive breast carcinoma (ATCC cat. HTB-22, RRID:CVCL_0031), MDA-MB-231 Breast adenocarcinoma, TNB (ATCC cat. HTB-26, RRID:CVCL_0062), as well as the small cell lung carcinoma cell lines NCI-H196 (ATCC cat. CRL-5823, CVCL_1509), NCI-H1417 (ATCC cat. CRL-5869, CVCL_1469), NCI-H1876 (ATCC cat. CRL-5902, CVCL_1503), and NCI-H510A (ATCC cat. HTB-184, CVCL_1565). The high grade ovarian serous adenocarcinoma cell lines OVCAR-5 (CVCL_1628) and OVCAR-8 (CVCL_1629) were established at the NCI and grown from internal seed stocks (33). The established alveolar rhabdomyosarcoma Rh18 (CVCL_1659) was provided by Dr. Peter Houghton (Greehey Children’s Cancer Research Institute, University of Texas Health Science Center, San Antonio, TX) (34,35). A list of the PDC and established cell lines is shown in **Table 2**. Pooled donor human umbilical vein endothelial cells (HUVEC, Lonza, cat. CC-2519) and human mesenchymal stem cells (hMSC, Lonza, cat. PT-2501) were purchased from Lonza (Walkersville, MD).

**Table 2.** Malignant cell lines grown as multi-cell type tumor spheroids (including endothelial cells and mesenchymal stem cells). The name, disease type, genetic variants, and the TP53 status are shown for each cell line.

| Cell Line Name | Disease Type | Genetic Variants | TP53 status |
| --- | --- | --- | --- |
| MDA-MB-231 | Breast adenocarcinoma, TNB | KRAS G13D, BRAF G464V, NF1 T467Hfs*3, NF2 E231*, EPHA3 E839*, CMTR2 C427Lfs*2 | R280K |
| 171881-019-R-J1 | Invasive breast carcinoma | PIK3CA E545Q, PIK3CA H1047R, MLH1 R265H | E271K |
| MCF7 | Invasive breast carcinoma | PIK3CA E545K, NBN R43*, EP300 R1356*, GATA3 D335Gfs*17, PTPRT X385 splice, PTPRD Y796* | WT |
| 885512-296-R-J2 | Invasive breast carcinoma, TNB | STK11 G251D, KMT2D Q3845*, RB1 S565*, ZNF750 P179Lfs*199 | I195T |
| 186277-243-T-J2 | Colon adenocarcinoma | KRAS G12D, DNMT3A R882H, PIK3CA C420R, CTCF R377C, ATM K2811Sfs*46 | E2D |
| 439559-082-T-J2 | Colon adenocarcinoma | KRAS G12D, APC R1450*, APC E941*, SOX9 Q391Pfs*13, AMER1 Q4Kfs*49 | R282W |
| 463943-066-T-J2 | Colon adenocarcinoma | KRAS G12V, APC T1493Rfs*14, APC S1068*, CDKN1B G97Vfs*22, EPHB1 R351Q, TCF7L2 S82Vfs*26, SMAD3 *426Yext*6, SOX9 P241Qfs*10 | WT |
| 519858-162-T-J1 | Colon adenocarcinoma | KRAS G12V, FLCN H429Pfs*27, APC T1023Nfs*6 | L323Tfs*14 |
| 616215-338-R-J1 | Colon adenocarcinoma | BRAF V600E, KRAS G12S, SMAD4 S32*, BRCA2 L1390Ffs*13, ATR I774Yfs*5, CDKN2A A4 P11dup | WT |
| OVCAR-5 | High grade ovarian serous adenocarcinoma | KRAS G12V, (PBRM1 X1526_splice, FANCD2 X126_splice, EP300 C1782* | WT |
| OVCAR-8 | High grade ovarian serous adenocarcinoma | ERBB2 G776V, CREBBP S893*, EP300 S1730Lfs*4, ATM V613L | c.376-1G>A |
| 556581-035-R-J1 | Ovarian clear cell carcinoma | PIK3CA Q546R, ARID1A G1282*, ARID1B Y1346Ifs*102, CTNNA1 D757Hfs*41 | WT |
| 323965-272-R-J2 | Pancreatic cancer (not Islets) | KRAS G12C, CDKN2A L16Pfs*9, ARID1A S958Lfs*10, PTPRS R1821* | R282W |
| 521955-158-R2-J5 | Pancreatic ductal adenocarcinoma | KRAS G12D, ATM G2287A | R158Sfs*8 |
| 521955-158-R6-J3 | Pancreatic ductal adenocarcinoma | KRAS G12D, ATM G2287A | R158Sfs*8 |
| PANC-1 | Pancreatic ductal adenocarcinoma | KRAS G12D | R273H |
| Rh18 | Alveolar rhabdomyosarcoma | PAX3-FOXO1 gene fusion | WT |
| 317291-083-R-J1 | Malignant fibrous histiocyoma | LATS1 C814Vfs*22 | WT |
| 596521-263-R-J1 | Malignant fibrous histiocyoma |  | L43* |
| MPNST | Malignant peripheral nerve sheath tumor | NF1 R304* | NA |
| 755483-174-R-J1 | Rhabdomyosarcoma | PBRM1 R1160*, LZTR1 X217_splice | WT |
| 772245-204-R-J1 | Rhabdomyosarcoma | PALB2 L939W | WT |
| 126254-015-R-J1 | Soft tissue sarcoma | RB1 E428Ifs*3 | WT |
| 494315-158-R-J1 | Soft tissue sarcoma | ZFP36L1 L198Sfs*35, ATRX S567* | WT |
| HS-SY-2 | Synovial sarcoma | SYT-SSX fusion | WT |
| SYO-1 | Synovial sarcoma | CTNNB1 G34L, SYT-SSX fusion, BRCA2 G2044V | WT |
| 327498-153-R-J2 | Uterine<br>carcinosarcoma | KRAS G12C, PTEN P38T, BCOR N1459S, CCND1 T286A,<br>ARID1A D1850Gfs*4, ARID1A P224Rfs*8, PTEN V45Ifs*6 | WT |
| 993429-296-R-J1 | Uterine<br>carcinosarcoma | PIK3CA E545K, NRAS G12D, RB1 E51*, PTEN V217G | X33_splice |
| NCI-H1417 | Small cell lung<br>carcinoma | RB1 Y321*, B2M L15Ffs*41, etoposide mid-response | R175L |
| NCI-H1876 | Small cell lung<br>carcinoma | KMT2D R5448*, CREBBP K1086*, NRAS Q61K, etoposide very<br>sensitive | R273L |
| NCI-H196 | Small cell lung<br>carcinoma | RB1 Q383*, PTEN Y138C, MED12 D395Efs*16, etoposide resistant | R175H |
| NCI-H510A | Small cell lung<br>carcinoma | RB1 K265*, SMAD4 G30*, BCORL1 E1172D, etoposide sensitive | R282G |

NA = Data not available
All data were obtained from the NCI Patient-Derived Models Repository (https://pdmr.cancer.gov/), cBioPortal (https://www.cbioportal.org/) Cellosaurus (https://www.cellosaurus.org/) and COSMIC (https://cancer.sanger.ac.uk/cell_lines).

### Cell Culture

All cells were maintained in an incubator at 37 °C and 5% CO_2_ with 95% humidity. The PDC lines were cultured according to standard operating procedures established by the NCI PDMR (https://pdmr.cancer.gov/sops/default.htm). Briefly, all PDCs were thawed and cultured in Matrigel-coated flasks prepared with a working solution of 1X Ham’s F-12 nutrient mix, without supplementation (Thermo Fisher Scientific, Waltham, MA, cat. 11765054), 100 U/mL penicillin-streptomycin (Thermo Fisher Scientific, cat. 15140122), and 2.5% Matrigel (Corning Inc., Corning, NY, cat. 354248) for the first three passages. All PDCs were cultured in complete DMEM/F-12 media containing advanced DMEM/F-12 (Thermo Fisher Scientific, cat. 12634028), 4.9% defined fetal bovine serum, heat inactivated (HyClone Laboratories Inc., Logan, UT, cat. SH30070.03HI), 389 ng/mL hydrocortisone (Sigma-Aldrich, cat. H4001), 9.7 ng/mL human EGF recombinant protein (Thermo Fisher Scientific, cat. PHG0313), 23.4 µg/mL adenine (Sigma-Aldrich, cat. A2786), 97.3 U/mL penicillin-streptomycin (Thermo Fisher Scientific, cat. 15140122), 1.9 mM L-glutamine (Thermo Fisher Scientific, cat. 25030081), and 9.7 µM Y-27632 dihydrochloride (Tocris Bioscience, Bristol, United Kingdom, cat. 1254). The PDCs were cultured in complete DMEM/F12 media without 9.7 µM Y-27632 dihydrochloride for at least two passages prior to the screen. The established cell lines HS-SY-2, SYO-1, and PANC-1 were cultured in DMEM, high glucose (Thermo Fisher Scientific, cat. 11965118) with 10% defined fetal bovine serum (HyClone Laboratories Inc., cat. SH30070.03). The established cell lines Rh18, OVCAR-5, OVCAR-8, MPNST, MCF7, MDA-MB-231, NCI-H196, NCI-H510A, and NCI-H1417 were cultured in RPMI 1640 medium, HEPES (Thermo Fisher Scientific, cat. 22400105) with 10% defined fetal bovine serum (HyClone Laboratories Inc., cat. SH30070.03). NCI-H1876 was cultured in DMEM/F-12, HEPES (Thermo Fisher Scientific, cat. 11330032), 5% defined fetal bovine serum (HyClone Laboratories Inc., cat. SH30070.03), ITS premix universal culture supplement [insulin (5 µg/mL), transferrin (5 µg/mL), and selenious acid (5 ng/mL)] (Corning Inc., cat. 354350), 10 nM hydrocortisone (Sigma-Aldrich, cat. H6909), 10 nM β-estradiol (Sigma-Aldrich, cat. E2257), and 4.5 mM L-glutamine (Thermo Fisher Scientific, cat. 25030081). The pooled donor HUVEC were cultured in endothelial cell growth medium 2 (PromoCell, Heidelberg, Germany, cat. C-22011), while the hMSC were cultured in mesenchymal stem cell growth medium 2 (PromoCell, cat. C-28009). For all experiments, HUVEC and hMSC were used at passages ≤5, while the malignant cell lines were used at passages ≤15. Samples of the cell lines were collected at regular intervals throughout the screening process for short tandem repeat (STR) profiling and mycoplasma testing by Labcorp (Laboratory Corporation of America Holdings, Burlington, NC, formerly known as Genetica DNA Laboratories) to confirm their authenticity and integrity.

### High-throughput Drug Combination Screening

Prior to their inoculation into microplates, malignant cells, HUVEC, and hMSC were removed from T flasks using TrypLE express (Thermo Fisher Scientific, cat. 12605036) and harvested by centrifugation for 5 min at 233 × g. Following removal of the supernatant, the cells were resuspended in fresh medium and counted using a Cellometer auto T4 bright field cell counter (Nexcelom, Lawrence, MA) and trypan blue to distinguish viable cells. Multi-cell type tumor spheroids were grown from the mixture of three cell types: 60% malignant cells, 25% HUVEC, and 15% hMSC (**Table S1**) as described previously (36,37). Mixed cell suspensions of 50 µL were dispensed into the wells of 384-well black/clear round-bottom ULA spheroid microplates (Corning Inc., cat. 3830). Following inoculation, the microplates were transferred to an incubator (Thermo Fisher Scientific) and maintained at 37 °C and 5% CO_2_ with 95% humidity. Three days after inoculation, test agents or controls were delivered to the wells of microplates. The approved and investigational anticancer agents, prepared as 800× stock solutions, were subsequently transferred in 62.5 nL volumes to the appropriate wells of microplates using an I.DOT non-contact dispenser (DISPENDIX, Stuttgart, Germany) to achieve a 1x final concentration. All anticancer agents and their combinations were tested in triplicate or quadruplicate. Additionally, each microplate included a DMSO vehicle control (*n* = 16) and a cytotoxicity control (1 µM staurosporine and 3 µM gemcitabine, *n* = 20). After delivery of the test agents and controls, the microplates were returned to the incubator for 7 days. Ten days after inoculation, the assay was completed with the addition of 20 µL CellTiter-Glo 3D (Promega, Madison, WI, cat. G9683) to each well. Next, the microplates were placed on a microplate shaker for 5 min. After 25 min of incubation at room temperature, luminescence was measured as a surrogate indicator of cell viability using a PHERAstar FSX microplate reader (BMG LABTECH, Cary, NC).

### Data Analysis

Luminescence measurements from the screen were exported as comma separated values (CSV) files and imported into custom Excel spreadsheets (Microsoft, Redmond, WA) for analysis. The raw luminescence data were evaluated for quality control, filtered for outliers, and converted to percent viability by normalizing to the DMSO (vehicle-treated) control. Concentration-response data were fit to the four-parameter logistic equation using the Solver Add-In in Excel. The Bliss independence model states that if two drugs have independent activities, then the viability for the combination is equal to the product of the viability measurements from the two single agents (38). Synergy between two compounds is indicated by a lower observed viability than predicted by the Bliss independence model, whereas antagonism is indicated by a greater observed viability than predicted.

### Data Availability

All data are accessible via the PubChem BioAssay public database (https://pubchem.ncbi.nlm.nih.gov/): AID 1963429; AID 1963428; AID 1963426; AID 1963427; AID 1963425; AID 1963424; AID 1963423; AID 1963422; AID 1963420; AID 1963421; AID 1963430; AID 1963451; AID 1963450; AID 1963449; AID 1963448; AID 1963434; AID 1963433; AID 1963447; AID 1963446; AID 1963444; AID 1963445; AID 1963443; AID 1963442; AID 1963440; AID 1963441; AID 1963439; AID 1963438; AID 1963436; AID 1963437; AID 1963435; AID 1963431; AID 1963432 (**Table S2**). The patient-derived cancer (PDC) cell line data used in this study can be downloaded from https://pdmr.cancer.gov/.

## Results

The three apoptosis pathway targeted agents alrizomadlin (APG-115), pelcitoclax (APG-1252), and dasminapant (APG-1387) were previously tested in the NCI-60 Human Tumor Cell Lines Screen (39) and their chemical structures are shown in **Figure 1** (**A – C, *top***). A COMPARE analysis (40) of the NCI-60 data indicated strong associations between the activity profile of each apoptosis pathway targeted agent and similar targeted agents with a Pearson correlation of >0.75. A force graph shows that alrizomadlin clustered with fourteen other MDM2 inhibitors, including idasanutlin (**Figure 1A, *middle***). Similarly, pelcitoclax clustered with eleven other Bcl-2/Bcl-xL inhibitors (**Figure 1B, *middle***), while dasminapant clustered with nine IAP family inhibitors (**Figure 1C, *middle***). Although alrizomadlin, pelcitoclax, and dasminapant all modulate cell death pathways, these results demonstrate their different activity profiles in cells that can be attributed to their distinct mechanisms of action.

**Figure 1.**
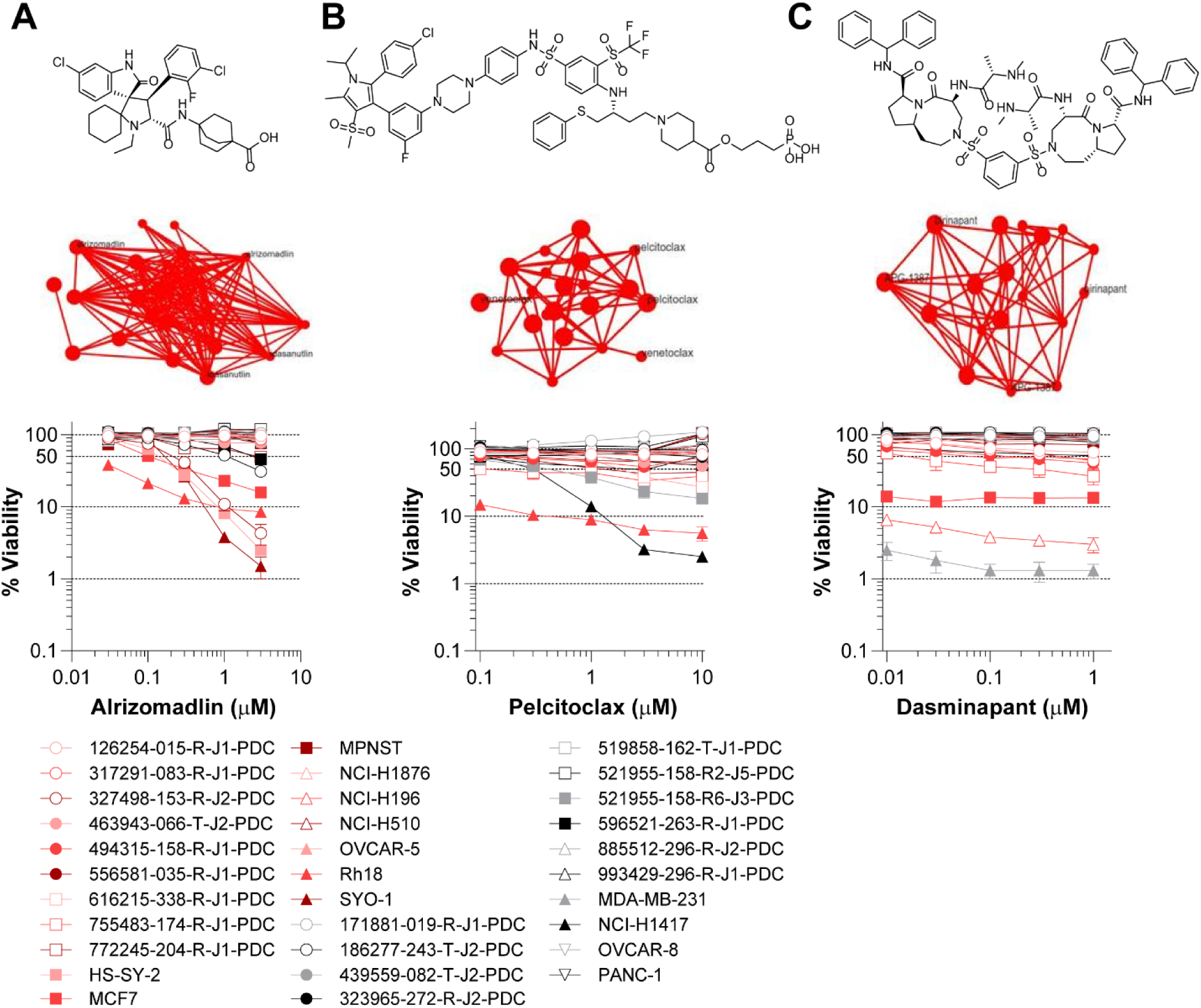
Single agent activity profiles of the apoptosis targeted agents alrizomadlin, pelcitoclax, and dasminapant. Chemical structures are shown for (A, *top*) alrizomadlin (APG-115), (B, *top*) pelcitoclax (APG-1252), and (C, *top*) dasminapant (APG-1387). Each agent was tested in the NCI-60 Human Tumor Cell Lines Screen and a COMPARE analysis was performed. The analysis demonstrated strong associations between each apoptosis pathway targeted agent and similar agents with a Pearson correlation of >0.75. Alrizomadlin (A, *middle*) clustered with fourteen MDM2 inhibitors, pelcitoclax (B, *middle*) clustered with eleven Bcl-2/Bcl-xL inhibitors and dasminapant (C, *middle*) clustered with nine IAP family inhibitors. A concentration-response graph with the data from all thirty-two multi-cell type tumor spheroid models is shown for (A, *bottom*) alrizomadlin, (B, *bottom*) pelcitoclax, and (C, *bottom*) dasminapant. Red symbols signify malignant cell lines with wildtype TP53, while gray symbols denote cell lines with a TP53 alteration (**Table 2**). All data show the mean ± SD (*n* ≥ 3 technical replicates).

To model the tumor microenvironment and its multicellular structure, multi-cell type tumor spheroids were utilized, which are amenable to high-throughput screening. This model incorporates both human malignant cells and non-malignant cells, including human umbilical vein endothelial cells (HUVEC) and human mesenchymal stem cells (hMSC) (36,37). The malignant cells were either established or patient-derived cell lines (**Table 2**). Before drug combinations were investigated (**Table 1**), the cytotoxicity of individual agents was evaluated across all thirty-two multi-cell type tumor spheroid models. The MDM2 inhibitor alrizomadlin demonstrated moderate selectivity for malignant cells carrying wildtype TP53, as evidenced by the concentration-response graphs (**Figure 1A, *bottom***) as well as area under the concentration-response curve (AUC) (**Figure S1A**). By AUC, the most sensitive tumor spheroids were the synovial sarcoma models, SYO-1 and HS-SY-2, as well as the alveolar rhabdomyosarcoma model, Rh18. The Bcl-2/Bcl-xL inhibitor pelcitoclax demonstrated minimal activity and only reached an IC_50_ in a limited number of spheroid models. The most sensitive models were the small cell lung carcinoma NCI-H1417 and the alveolar rhabdomyosarcoma Rh18 (**Figure 1B, *bottom*** and **Figure S1B**). Similarly, the IAP antagonist and Smac mimetic dasminapant displayed minimal cytotoxicity and only reached an IC_50_ in several spheroid models (**Figure 1C, *bottom*** and **Figure S1C**). The invasive breast carcinoma model MCF7, the triple-negative breast adenocarcinoma model MDA-MB-231, and the small cell lung carcinoma model NCI-H196 exhibited the greatest sensitivity to dasminapant as a single agent.

Combinations with the apoptosis targeted agents were assessed by the Bliss independence model (38). Within this model, an observed combination response is juxtaposed with the predicted combination response, under the assumption that the agents act independently and do not interact with each other. Bliss independence scores near zero signify an additive effect, where the observed response closely matches the predicted outcome. Positive Bliss scores imply greater-than-additive effects or synergy, whereas negative Bliss scores indicate antagonism. A mean Bliss matrix score was calculated for each spheroid model from each combination’s concentration matrix ([5 concentrations of drug A, *n* ≥ 3 technical replicates] × [6 concentrations of drug B, *n* ≥ 3 technical replicates] = [30 combination concentrations]). A comprehensive summary of the mean Bliss scores (*n* = 1,280) from all combinations tested (*n* = 40) in all spheroid models (*n* = 32) is shown in **Figure S2** and **Table S3**.

Among the most active combinations was alrizomadlin and pelcitoclax, which enabled dual targeting of the apoptosis pathway. **Figure 2A** shows concentration-response data and Bliss independence scores across the concentration matrices from six selected tumor spheroid models, including four with a wildtype TP53 status, one with a E2D variant, and one with a R175L variant (**Table 2**). As single agents, alrizomadlin and pelcitoclax exhibited minimal activity in the 186277-243-T-J2 and 616215-338-R-J1 colon adenocarcinoma models, as well as the 772245-204-R-J1 rhabdomyosarcoma model; however, their combination demonstrated synergy. The SYO-1 and HS-SY-2 synovial sarcoma models were insensitive to pelcitoclax but highly responsive to alrizomadlin. The addition of pelcitoclax potentiated the cytotoxicity of alrizomadlin. Conversely, in the small cell lung carcinoma model NCI-H1417, with a TP53 R175L variant, alrizomadlin potentiated the cytotoxicity of pelcitoclax. Combination effects were also observed between alrizomadlin and dasminapant, albeit to a lesser extent. **Figure 2B** shows selected data from three sarcoma models, wherein dasminapant enhanced the cytotoxicity of alrizomadlin.

**Figure 2.**
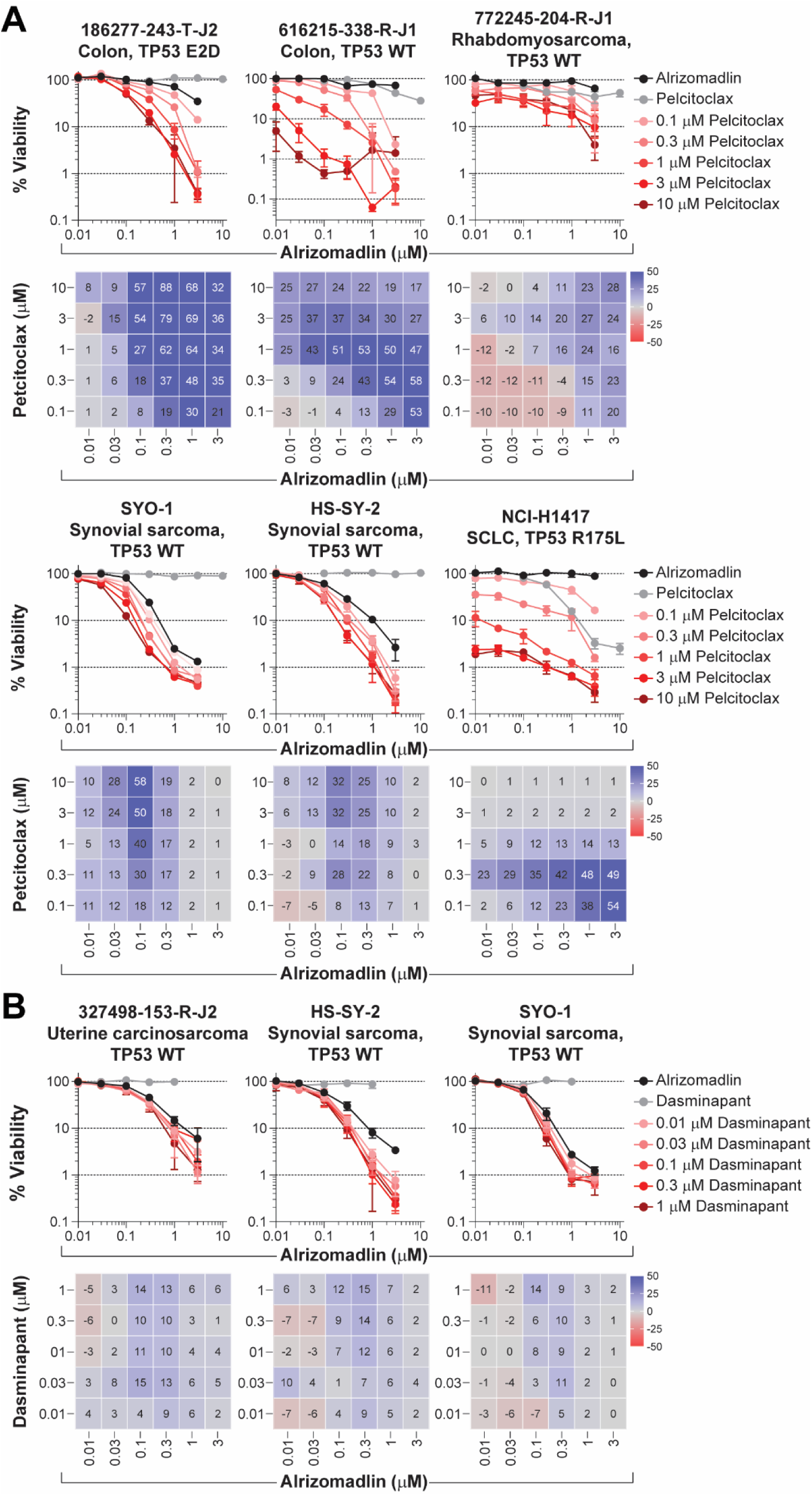
Dual apoptosis pathway targeting by the combination of alrizomadlin with pelcitoclax or dasminapant. Concentration-response graphs (*top*, mean ± SD, *n* ≥ 3 technical replicates) from combinations of alrizomadlin with A) pelcitoclax or B) dasminapant are shown with corresponding Bliss independence scores from each combination’s concentration matrix (*bottom*, mean of *n* ≥ 3 technical replicates) displayed numerically and as a heat map (blue indicates synergy, gray indicates additivity, and red indicates antagonism). The malignant cell line name, disease type, and TP53 status are indicated above each graph.

Alrizomadlin in combination with the XPO1 (exportin-1) inhibitor eltanexor demonstrated additive to greater-than additive cytotoxicity. **Figure 3A** shows concentration-response data and Bliss independence scores across the concentration matrices from four selected multi-cell type tumor spheroid models treated with alrizomadlin and eltanexor. In three tumor spheroid models alrizomadlin demonstrated synergistic activity with eltanexor across various concentrations. Although eltanexor showed no single-agent activity in the 616215-338-R-J1 colon adenocarcinoma model, substantial cytotoxicity was observed in combination with alrizomadlin. In all four models, this combination resulted in >99% cytotoxicity (two-log reduction). Similar outcomes were noted when eltanexor was combined with pelcitoclax (**Figure 3B**). As single agents, pelcitoclax demonstrated no activity in the 186277-243-T-J2 colon adenocarcinoma model, while alrizomadlin exhibited no activity in the 616215-338-R-J1 colon adenocarcinoma model. However, their combination demonstrated substantial synergistic activities in these models. Additionally, the combination of pelcitoclax and eltanexor demonstrated synergy in both the 772245-204-R-J1 rhabdomyosarcoma and NCI-H1417 small cell lung carcinoma models.

**Figure 3.**
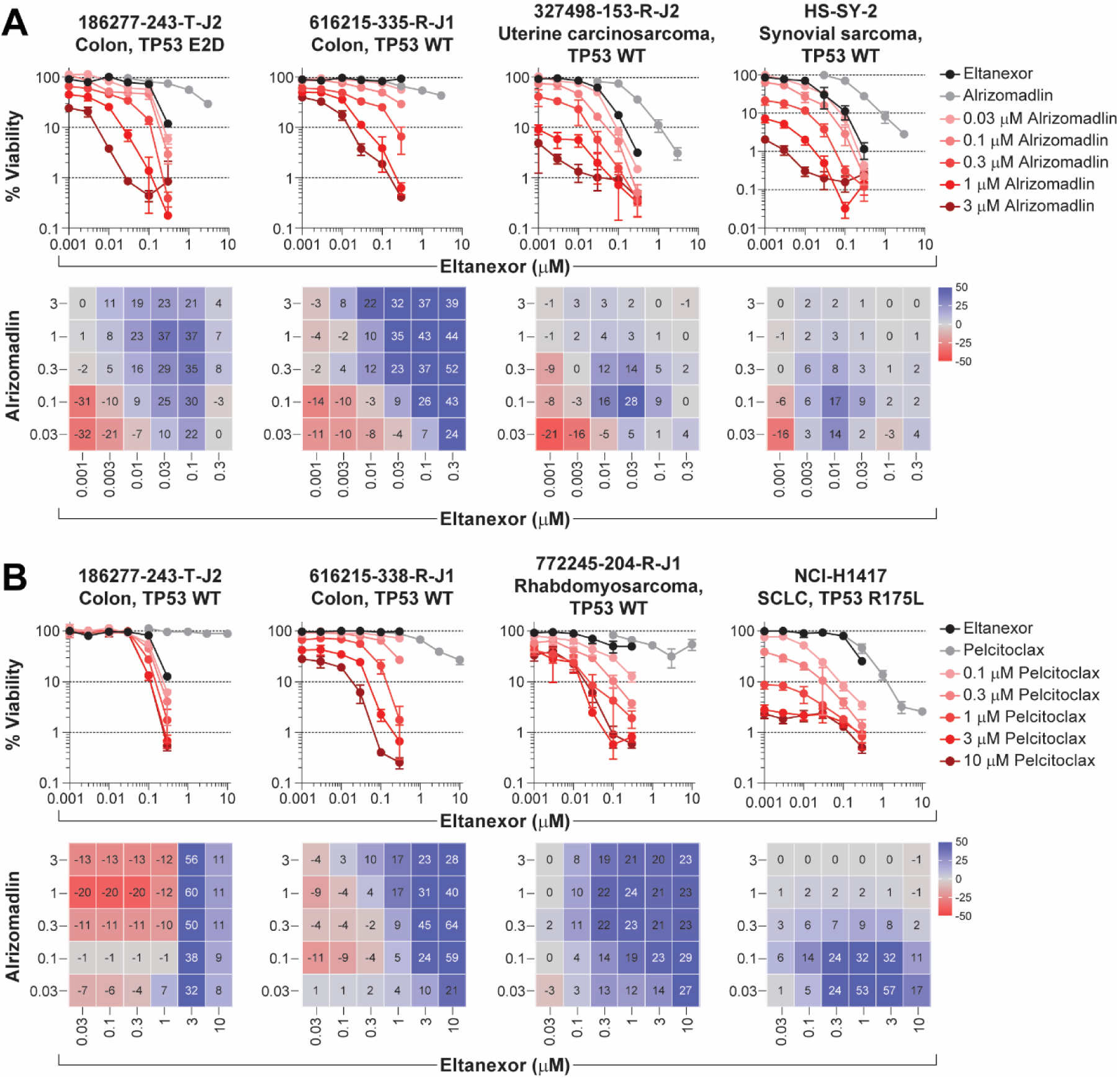
Combination of the XPO1 inhibitor eltanexor with the apoptosis targeted agent alrizomadlin or pelcitoclax. Concentration-response graphs (*top*, mean ± SD, *n* ≥ 3 technical replicates) from combinations of eltanexor with A) alrizomadlin or B) pelcitoclax are shown with corresponding Bliss independence scores from each combination’s concentration matrix (*bottom*, mean of *n* ≥ 3 technical replicates) displayed numerically and as a heat map (blue indicates synergy, gray indicates additivity, and red indicates antagonism). The malignant cell line name, disease type, and TP53 status are indicated above each graph.

Another combination strategy was to pair the apoptosis targeted agents with the pan-PI3K inhibitor copanlisib. **Figure 4A** shows concentration-response data and Bliss independence scores across the concentration matrices from the combination of alrizomadlin and copanlisib in selected multi-cell type tumor spheroid models. In all four models, this combination demonstrated additive to greater-than-additive activity and induced >99% cytotoxicity. Likewise, pelcitoclax also demonstrated additive and synergistic effects in combination with copanlisib, as observed from four selected multi-cell type tumor spheroid models shown in **Figure 4B**. Notably, synergistic effects were evident in the 616215-338-R-J1 colon adenocarcinoma and NCI-H1417 small cell lung carcinoma models. Pelcitoclax did not show single-agent activity in the 186277-243-T-J2 colon adenocarcinoma or the NCI-H1876 small cell lung carcinoma models. However, the cytotoxicity of copanlisib at higher concentrations was potentiated by pelcitoclax. PARP inhibitors, which target DNA repair pathways, also demonstrated synergy with pelcitoclax. **Figure 5** shows concentration-response graphs and Bliss independence scores across the concentration matrices from pelcitoclax in combination with the PARP inhibitors olaparib (**Figure 5A**) and talazoparib (**Figure 5B**) in four selected multi-cell type tumor spheroid models. Similarities in response were noted for olaparib and talazoparib within the same tumor model.

**Figure 4.**
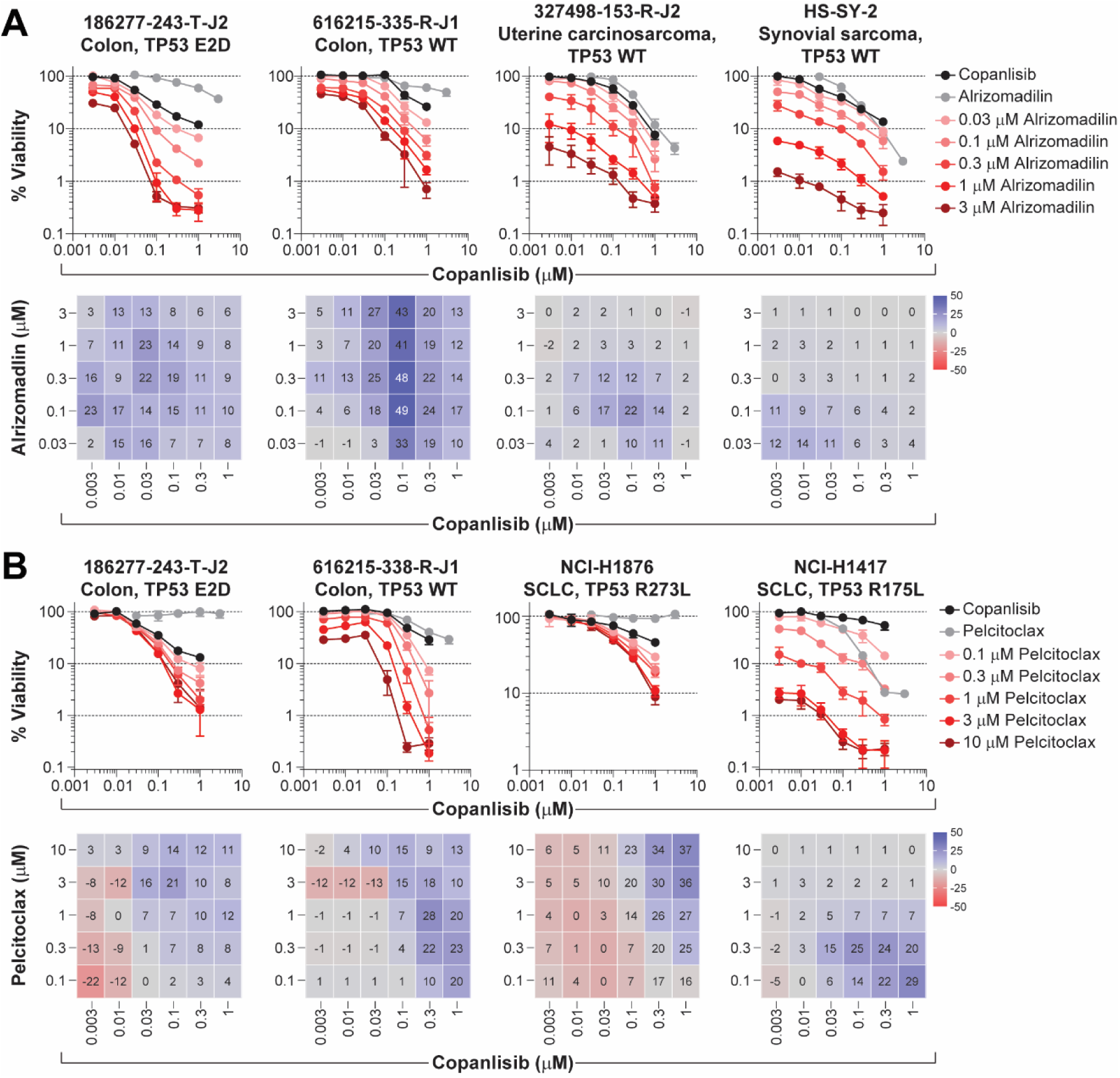
Combination of the pan-PI3K inhibitor copanlisib with the apoptosis targeted agent alrizomadlin or pelcitoclax. Concentration-response graphs (*top*, mean ± SD, *n* ≥ 3 technical replicates) from combinations of copanlisib with A) alrizomadlin or B) pelcitoclax are shown with corresponding Bliss independence scores from each combination’s concentration matrix (*bottom*, mean of *n* ≥ 3 technical replicates) displayed numerically and as a heat map (blue indicates synergy, gray indicates additivity, and red indicates antagonism). The malignant cell line name, disease type, and TP53 status are indicated above each graph.

**Figure 5.**
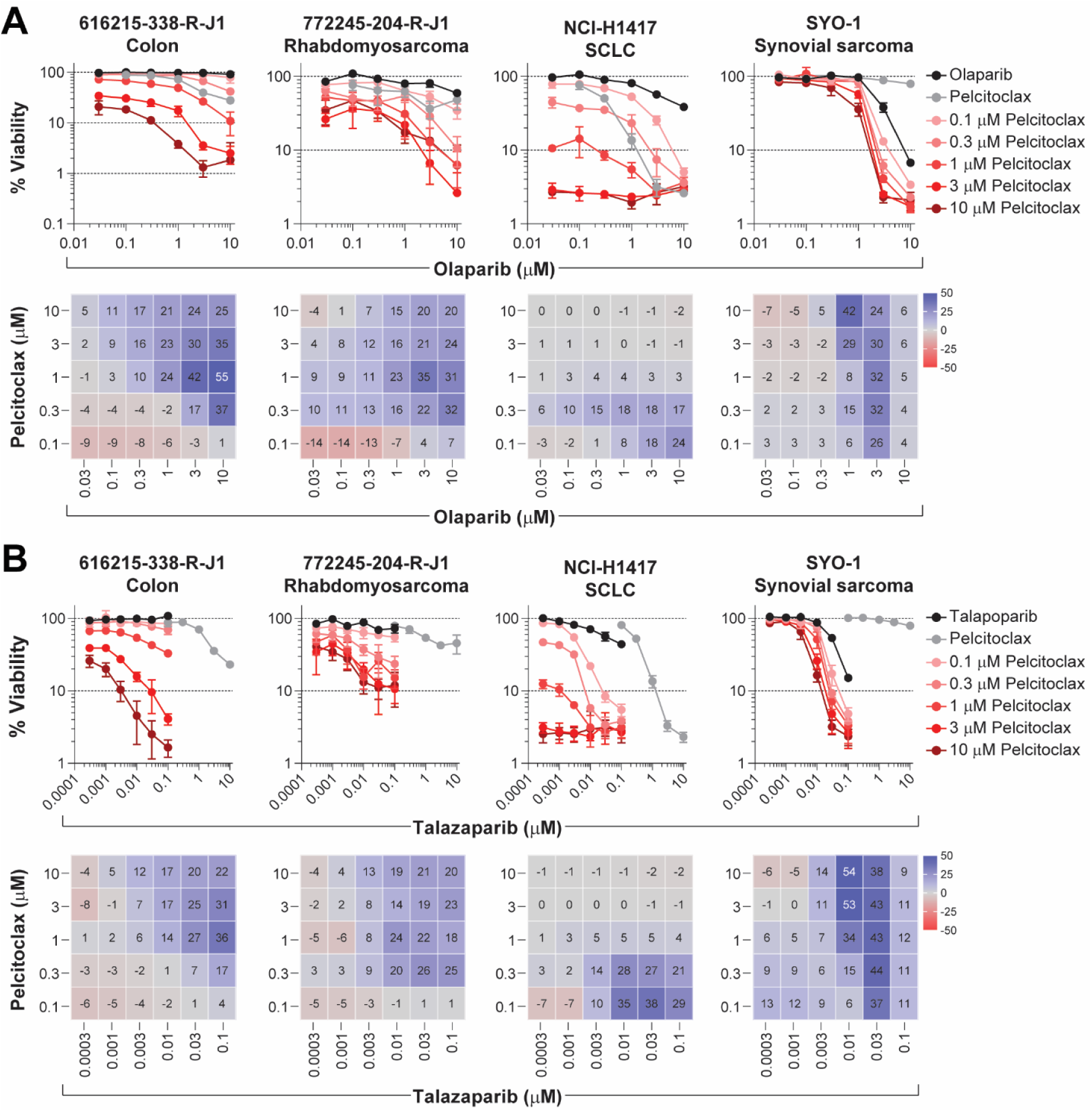
Combination of the apoptosis targeted agent pelcitoclax with the PARP inhibitor olaparib or talazoparib. Concentration-response graphs (*top*, mean ± SD, *n* ≥ 3 technical replicates) from combinations of pelcitoclax with A) olaparib or B) talazoparib are shown with corresponding Bliss independence scores from each combination’s concentration matrix (*bottom*, mean of *n* ≥ 3 technical replicates) displayed numerically and as a heat map (blue indicates synergy, gray indicates additivity, and red indicates antagonism). The malignant cell line name and disease type are indicated above each graph.

Three inhibitors that target the MAPK pathway were also evaluated, including two KRAS variant specific inhibitors, MRTX1133 and adagrasib, and the MEK inhibitor cobimetinib. **Figure 6A** shows concentration-response graphs and Bliss independence scores across the concentration matrices for the KRAS G12D inhibitor MRTX1133 in combination with each of the apoptosis pathway targeted agents. All data shown are from the 186277-243-T-J2 colon adenocarcinoma model with a KRAS G12D variant. Alrizomadlin demonstrated both additivity and synergy with MRTX1133 and the combination achieved approximately 99% cytotoxicity at the highest concentrations. Pelcitoclax and dasminapant, while inactive as single agents, potentiated the cytotoxicity of MRTX1133 when combined. In the uterine carcinosarcoma model 327498-153-R-J2, with a KRAS G12C variant, the KRAS G12C-specific inhibitor adagrasib showed combination effects with alrizomadlin, but not pelcitoclax or dasminapant (**Figure 6B**). Lastly, in the 616215-338-R-J1 colon adenocarcinoma model, which carries KRAS G12S and BRAF V600E variants, both alrizomadlin and pelcitoclax exhibited synergy in combination with the MEK1 inhibitor cobimetinib, whereas the effects were diminished with dasminapant (**Figure 6C**).

**Figure 6.**
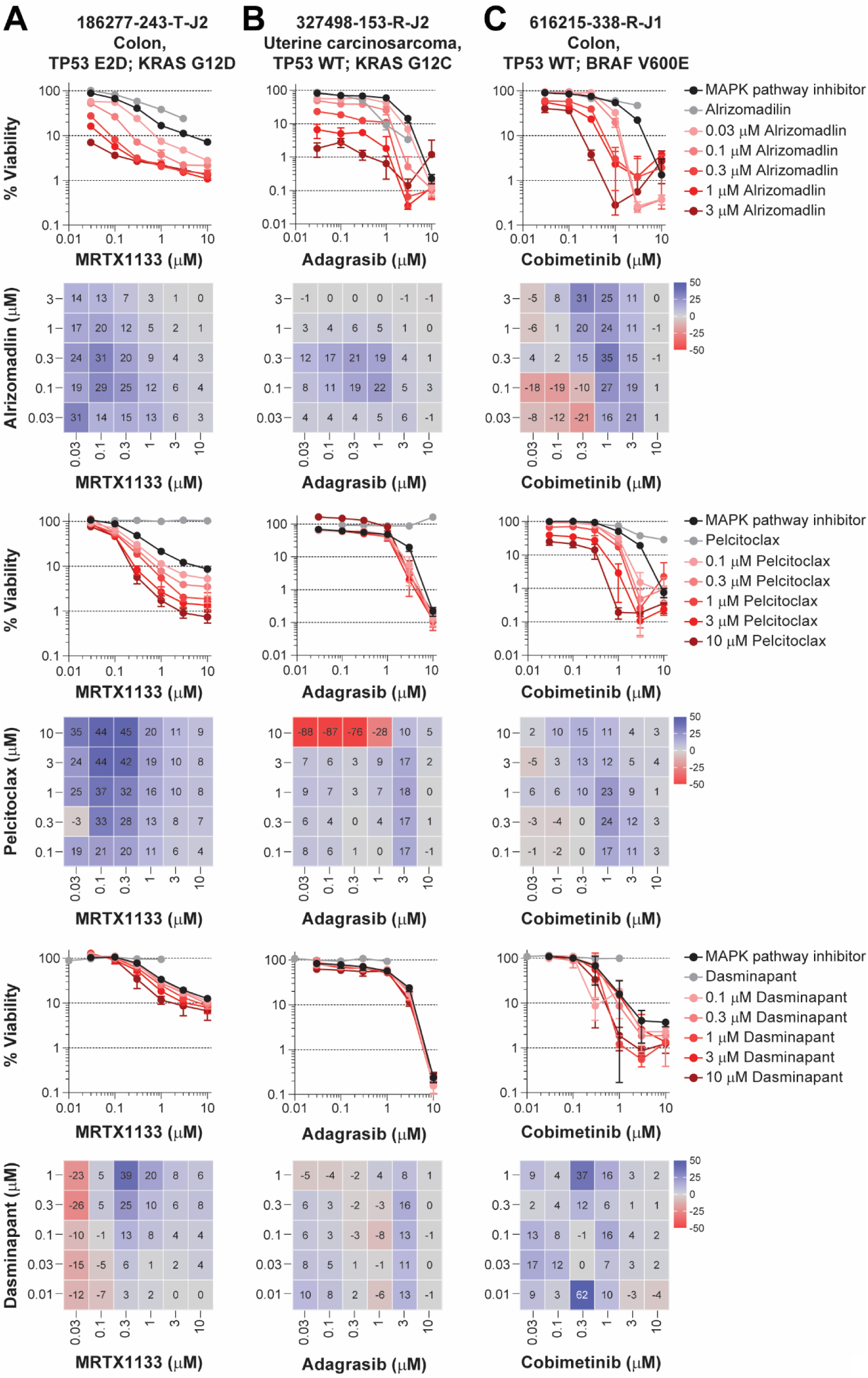
Combination of apoptosis targeted agents with MAPK pathway inhibitors. A) Concentration-response graphs (mean ± SD, *n* ≥ 3 technical replicates) from the KRAS G12D inhibitor MRTX1133 in combination with the apoptosis targeted agents (*top*, alrizomadlin; *middle*, pelcitoclax; *bottom*, dasminapant) in the 186277-243-T-J2 colon adenocarcinoma model harboring a KRAS G12D variant. B) Concentration-response graphs (mean ± SD, *n* ≥ 3 technical replicates) from the KRAS G12C inhibitor adagrasib in combination with the apoptosis targeted agents (*top*, alrizomadlin; *middle*, pelcitoclax; *bottom*, dasminapant) in the 327498-153-R-J2 uterine carcinosarcoma model harboring a KRAS G12C variant. C) Concentration-response graphs (mean ± SD, *n* ≥ 3 technical replicates) from the MEK inhibitor cobimetinib in combination with the apoptosis targeted agents (*top*, alrizomadlin; *middle*, pelcitoclax; *bottom*, dasminapant) in the 616215-338-R-J1 colon adenocarcinoma model harboring a BRAF V600E variant. Below each concentration-response graph, corresponding Bliss independence scores are shown from each combination’s concentration matrix (mean of *n* ≥ 3 technical replicates) displayed numerically and as a heat map (blue indicates synergy, gray indicates additivity, and red indicates antagonism). The malignant cell line name, disease type, TP53 status, and oncogenic variants are indicated above each graph.

## Discussion

Apoptosis is an essential regulatory mechanism in cells and dysregulation of the programmed cell death processes could result in the survival and proliferation of damaged cells destined to become malignant (1). An important segment of cancer drug development has focused on restoring apoptotic pathways in cancer cells. This study examined varied sarcomas and major solid tumor models after exposure to the MDM2 inhibitor alrizomadlin, the Bcl-2/Bcl-xL inhibitor pelcitoclax or the IAP protein family antagonist dasminapant in combination with standard of care or investigational small molecule anticancer agents.

Sensitivity of the thirty-two multi-cell type tumor spheroid models to the MDM2 inhibitor alrizomadlin was primarily limited to the synovial sarcoma models SYO-1 and HS-SY-2, the uterine carcinosarcoma model 327498-153-R-J2 and the alveolar rhabdomyosarcoma model Rh18. The invasive breast carcinoma model MCF7 also showed above average sensitivity to alrizomadlin (**Figure 1A, *bottom***). The activity of alrizomadlin against the SYO-1 and HS-SY-2 models was potentiated by both pelcitoclax (**Figure 2A**) and dasminapant (**Figure 2B**). Several models that were minimally sensitive to alrizomadlin as a single agent demonstrated substantial sensitivity to the combination of alrizomadlin and pelcitoclax, including the colon adenocarcinoma models 186277-243-T-J2 and 616215-338-R-J1, the rhabdomyosarcoma model 772245-204-R-J1, and the small cell lung carcinoma model NCI-H1417 (**Figure 2A**). It is noteworthy that the p53 pathway is altered in the 186277-243-T-J2, 616215-338-R-J1, and NCI-H1417 models. The 186277-243-T-J2 contains a TP53 E2D variant and ATM K2811Sfs*46 alteration, while the 616215-338-R-J1 model has a CDKN2A A4_P11dup alteration. The NCI-H1417 model contains a R175L variant of TP53 (**Table 2**). There are no apparent alterations to the p53 pathway of the 772245-204-R-J1 model, but its sensitivity to the combination was reduced compared to the other three models. Combinations of MDM2 and Bcl-2 inhibitors have previously been shown to induce synthetic lethality in resistant AML mouse models (41). In TP53 wildtype B-cell acute lymphoblastic leukemias (ALL), a combination of the MDM2 inhibitor idasanutlin and the Bcl-2 inhibitor venetoclax was explored in vitro and showed synergy in three ALL cell lines representing Ph+, Ph-, and Ph-like ALL (42). In solid tumors, synergistic effects via co-inhibition of MDM2 and Bcl-2/Bcl-xL were observed in neuroblastoma orthotopic xenograft tumors and uveal melanoma cell lines (43,44). Taken together, the results indicate that combination regimes based on alrizomadlin-mediated p53 reactivation are successful across many cancer types.

The combination of alrizomadlin and the XPO1 inhibitor eltanexor showed additive to greater-than-additive responses in the colon adenocarcinoma models 186277-243-T-J2 and 616215-338-R-J1, the uterine carcinosarcoma model 327498-153-R-J2, and the synovial sarcoma model HS-SY-2 (**Figure 3A**). While the colon models were minimally responsive to the single agents, the sarcoma models demonstrated substantial sensitivity to both. All four models were highly sensitive to the combination and demonstrated more than two logs of cytotoxicity. As noted above, the p53 pathway is altered in the 186277-243-T-J2 and 616215-338-R-J1 models; however, the pathway does not appear to be altered in the sarcoma models 327498-153-R-J2 and HS-SY-2. Additive and greater-than-additive activities were also observed in these four models from the combination of alrizomadlin and the pan-PI3K inhibitor copanlisib (**Figure 4A**). Moreover, the combination reduced cell viability of all four models by more than two logs.

The colon adenocarcinoma model 186277-243-T-J2 contains a KRAS G12D variant (**Table 2**) and the combination of alrizomadlin with the KRAS G12D-selective inhibitor MRTX1133 demonstrated additive and synergistic activity (**Figure 6A**). Similarly, the uterine carcinosarcoma model 327498-153-R-J2 harbors a KRAS G12C variant and the KRAS G12C-selective drug adagrasib demonstrated additive and greater-than-additive responses in combination with alrizomadlin (**Figure 6B**). Finally, alrizomadlin demonstrated synergy with the MEK inhibitor cobimetinib in the colon adenocarcinoma model 616215-338-R-J1, which contains KRAS G12S and BRAF V600E variants (**Figure 6C**). The responses in these models reached or exceeded two logs of cytotoxicity.

Pelcitoclax (APG-1252) is a novel dual inhibitor of Bcl-2 and Bcl-xL proteins that was designed as a prodrug to enhance its therapeutic index, lower cell permeability, and reduce platelet toxicity. After administration, pelcitoclax undergoes conversion into the more potent metabolite APG-1252-M1, which exhibits significantly higher levels in tumors compared to plasma. This mechanism accounts for the reduction in Bcl-xL-driven on-target thrombocytopenia (45). In this study, pelcitoclax demonstrated limited activity, as a single agent, among the thirty-two multi-cell type tumor spheroid models. The small cell lung carcinoma model NCI-H1417 and the alveolar rhabdomyosarcoma model Rh18 showed the greatest sensitivity to pelcitoclax (**Figure 1B, *bottom***). Small cell lung cancer has relatively high expression of Bcl-2 protein and tends to be more responsive than other solid tumors to Bcl-2 inhibitors (46). Human sarcoma cell lines were shown to be responsive to Bcl-2 inhibitors (35) and the demonstration of Bcl-2 addiction in synovial sarcoma points to novel therapeutic combination approaches with Bcl-2 protein family targeted agents (47,48). Similar to alrizomadlin, pelcitoclax demonstrated synergy with the XPO1 inhibitor eltanexor in the colon adenocarcinoma models 186277-243-T-J2 and 616215-338-R-J1 (**Figure 3B**). Preclinical studies have demonstrated activity of the XPO1 inhibitor selinexor in various sarcoma subtypes (49). The combination of pelcitoclax and eltanexor was highly synergistic in the 772245-204-R-J1 rhabdomyosarcoma model, which does not have any apparent alterations within the p53 pathway. The combination also showed additive and synergistic activity in the NCI-H1417 small cell lung carcinoma model. Several recent studies have reported synergy from combinations of agents that inhibit Bcl-2/Bcl-xL and XPO1 (50–52). In combination with copanlisib, pelcitoclax showed additive to greater-than-additive activity that was comparable to the combination of alrizomadlin and copanlisib (**Figure 4B**). A recent first-in-human study of pelcitoclax in combination with paclitaxel demonstrated modest activity in patients with relapsed/refractory small-cell lung cancer (53).

In combination with the PARP inhibitors olaparib and talazoparib, pelcitoclax demonstrated additive and synergistic activity, which reduced cell viability by more than one log in the four models shown in **Figure 5**. While the single agents showed modest activity in the 616215-338-R-J1 colon adenocarcinoma and 772245-204-R-J1 rhabdomyosarcoma models, the combinations increased cytotoxicity by approximately one log. Very little combination activity was observed between pelcitoclax and adagrasib in the 327498-153-R-J2 model with a KRAS G12C variant; however, substantial synergy was observed between pelcitoclax and MRTX1133 in the 186277-243-T-J2 model containing a KRAS G12D variant with cytotoxicity approaching two logs (**Figure 6**). Additive and greater-than-additive activities were also observed from the combination of pelcitoclax and cobimetinib in the 616215-338-R-J1 colon adenocarcinoma model with BRAF V600E and KRAS G12S variants.

Overall, the thirty-two multi-cell type tumor spheroid models demonstrated relatively modest and shallow concentration responses to the IAP protein family antagonist dasminapant as a single agent. The breast models MDA-MB-231 and MCF7 were most sensitive along with the small cell lung carcinoma model NCI-H196 (**Figure 1C, *bottom***). Compared to alrizomadlin and pelcitoclax, dasminapant showed similar, but overall reduced activity in combination with other agents. For example, in combination with alrizomadlin, pelcitoclax showed greater synergy in the synovial sarcoma models than dasminapant (**Figure 2**). Furthermore, MRTX1133, adagrasib, and cobimetinib showed fewer combination effects with dasminapant compared to alrizomadlin and pelcitoclax (**Figure 6**).

This study demonstrated a greater-than-additive effect with MDM2 and Bcl-2/Bcl-xL inhibitors, alrizomadlin and pelcitoclax, in a range of solid tumor multi-cell type tumor spheroid models including colon, breast, uterine sarcoma, rhabdomyosarcoma, synovial sarcoma, and SCLC, suggesting that this novel combination should be tested *in vivo*. These two agents are currently under phase 1b/2 clinical investigations, including combination with immunotherapy for alrizomadlin (NCT04785196) and small molecules for pelcitoclax (NCT04001777, NCT05691504). Due to some overlapping toxicity such as thrombocytopenia, which was reported in phase 1 studies (54,55), a combination trial of alrizomadlin and pelcitoclax could be conducted with caution by exploring different dosing schedules.

## References

1. Koren E, Fuchs Y. Modes of Regulated Cell Death in Cancer. Cancer Discov 2021;11:245–65

2. Neophytou CM, Trougakos IP, Erin N, Papageorgis P. Apoptosis Deregulation and the Development of Cancer Multi-Drug Resistance. Cancers (Basel) 2021;13

3. Carneiro BA, El-Deiry WS. Targeting apoptosis in cancer therapy. Nat Rev Clin Oncol 2020;17:395–417

4. Jan R, Chaudhry GE. Understanding Apoptosis and Apoptotic Pathways Targeted Cancer Therapeutics. Adv Pharm Bull 2019;9:205–18

5. Kandoth C, McLellan MD, Vandin F, Ye K, Niu B, Lu C, et al. Mutational landscape and significance across 12 major cancer types. Nature 2013;502:333–9

6. Hafner A, Bulyk ML, Jambhekar A, Lahav G. The multiple mechanisms that regulate p53 activity and cell fate. Nat Rev Mol Cell Biol 2019;20:199–210

7. Chen X, Zhang T, Su W, Dou Z, Zhao D, Jin X, et al. Mutant p53 in cancer: from molecular mechanism to therapeutic modulation. Cell Death Dis 2022;13:974

8. Weinberg RA. Coming full circle-from endless complexity to simplicity and back again. Cell 2014;157:267–71

9. Ventura A, Kirsch DG, McLaughlin ME, Tuveson DA, Grimm J, Lintault L, et al. Restoration of p53 function leads to tumour regression in vivo. Nature 2007;445:661–5

10. Xue W, Zender L, Miething C, Dickins RA, Hernando E, Krizhanovsky V, et al. Senescence and tumour clearance is triggered by p53 restoration in murine liver carcinomas. Nature 2007;445:656–60

11. Aguilar A, Lu J, Liu L, Du D, Bernard D, McEachern D, et al. Discovery of 4-((3’R,4’S,5’R)-6’’-Chloro-4’-(3-chloro-2-fluorophenyl)-1’-ethyl-2’’-oxodispiro[cyclohexane-1,2’-pyrrolidine-3’,3’’-indoline]-5’-carboxamido)bicyclo[2.2.2]octane-1-carboxylic Acid (AA-115/APG-115): A Potent and Orally Active Murine Double Minute 2 (MDM2) Inhibitor in Clinical Development. J Med Chem 2017;60:2819–39

12. Iksen, Witayateeraporn W, Hardianti B, Pongrakhananon V. Comprehensive review of Bcl-2 family proteins in cancer apoptosis: Therapeutic strategies and promising updates of natural bioactive compounds and small molecules. Phytother Res 2024

13. Zhang L, Lu Z, Zhao X. Targeting Bcl-2 for cancer therapy. Biochim Biophys Acta Rev Cancer 2021;1876:188569

14. Zhang Z, Bai L, Hou L, Deng H, Luan S, Liu D, et al. Trends in targeting Bcl-2 anti-apoptotic proteins for cancer treatment. Eur J Med Chem 2022;232:114184

15. Kaloni D, Diepstraten ST, Strasser A, Kelly GL. BCL-2 protein family: attractive targets for cancer therapy. Apoptosis 2023;28:20–38

16. Campbell KJ, Tait SWG. Targeting BCL-2 regulated apoptosis in cancer. Open Biol 2018;8

17. Diepstraten ST, Anderson MA, Czabotar PE, Lessene G, Strasser A, Kelly GL. The manipulation of apoptosis for cancer therapy using BH3-mimetic drugs. Nat Rev Cancer 2022;22:45–64

18. DiNardo CD, Pratz K, Pullarkat V, Jonas BA, Arellano M, Becker PS, et al. Venetoclax combined with decitabine or azacitidine in treatment-naive, elderly patients with acute myeloid leukemia. Blood 2019;133:7–17

19. Bai L, Chen J, Liu L, McEachern D, Aguilar A, Zhou H, et al. BM-1252 (APG-1252): a potent dual specific Bcl-2/Bcl-xL inhibitor that achieves complete tumor regression with minimal platelet toxicity. Eur J Cancer 2014;50:109–10

20. Zhang H, Nimmer PM, Tahir SK, Chen J, Fryer RM, Hahn KR, et al. Bcl-2 family proteins are essential for platelet survival. Cell Death Differ 2007;14:943–51

21. Wilson WH, O’Connor OA, Czuczman MS, LaCasce AS, Gerecitano JF, Leonard JP, et al. Navitoclax, a targeted high-affinity inhibitor of BCL-2, in lymphoid malignancies: a phase 1 dose-escalation study of safety, pharmacokinetics, pharmacodynamics, and antitumour activity. Lancet Oncol 2010;11:1149–59

22. Cetraro P, Plaza-Diaz J, MacKenzie A, Abadía-Molina F. A Review of the Current Impact of Inhibitors of Apoptosis Proteins and Their Repression in Cancer. Cancers 2022;14

23. Kumar S, Fairmichael C, Longley DB, Turkington RC. The Multiple Roles of the IAP Super-family in cancer. Pharmacol Therapeut 2020;214

24. Wang SM. Design of Small-Molecule Smac Mimetics as IAP Antagonists. Curr Top Microbiol 2011;348:89–113

25. Zhao XY, Wang XY, Wei QY, Xu YM, Lau ATY. Potency and Selectivity of SMAC/DIABLO Mimetics in Solid Tumor Therapy. Cells-Basel 2020;9

26. Cossu F, Milani M, Mastrangelo E, Lecis D. Targeting the BIR Domains of Inhibitor of Apoptosis (IAP) Proteins in Cancer Treatment. Comput Struct Biotec 2019;17:142–50

27. Cerna D, Lim B, Adelabu Y, Yoo S, Carter D, Fahim A, et al. SMAC Mimetic/IAP Inhibitor Birinapant Enhances Radiosensitivity of Glioblastoma Multiforme. Radiat Res 2021;195:549–60

28. Srivastava AK, Jaganathan S, Stephen L, Hollingshead MG, Layhee A, Damour E, et al. Effect of a Smac Mimetic (TL32711, Birinapant) on the Apoptotic Program and Apoptosis Biomarkers Examined with Validated Multiplex Immunoassays Fit for Clinical Use. Clin Cancer Res 2016;22:1000–10

29. Chen Z, Chen J, Liu H, Dong W, Huang X, Yang D, et al. The SMAC Mimetic APG-1387 Sensitizes Immune-Mediated Cell Apoptosis in Hepatocellular Carcinoma. Front Pharmacol 2018;9:1298

30. Li BX, Wang HB, Qiu MZ, Luo QY, Yi HJ, Yan XL, et al. Correction to: Novel smac mimetic APG-1387 elicits ovarian cancer cell killing through TNF-alpha, Ripoptosome and autophagy mediated cell death pathway. J Exp Clin Cancer Res 2018;37:108

31. Li N, Feng L, Han HQ, Yuan J, Qi XK, Lian YF, et al. A novel Smac mimetic APG-1387 demonstrates potent antitumor activity in nasopharyngeal carcinoma cells by inducing apoptosis. Cancer Lett 2016;381:14–22

32. Pan W, Luo Q, Yan X, Yuan L, Yi H, Zhang L, et al. A novel SMAC mimetic APG-1387 exhibits dual antitumor effect on HBV-positive hepatocellular carcinoma with high expression of cIAP2 by inducing apoptosis and enhancing innate anti-tumor immunity. Biochem Pharmacol 2018;154:127–35

33. Schilder RJ, Hall L, Monks A, Handel LM, Fornace AJ, Jr., Ozols RF, et al. Metallothionein gene expression and resistance to cisplatin in human ovarian cancer. Int J Cancer 1990;45:416–22

34. Hazelton BJ, Houghton JA, Parham DM, Douglass EC, Torrance PM, Holt H, Houghton PJ. Characterization of cell lines derived from xenografts of childhood rhabdomyosarcoma. Cancer Res 1987;47:4501–7

35. Teicher BA, Polley E, Kunkel M, Evans D, Silvers T, Delosh R, et al. Sarcoma Cell Line Screen of Oncology Drugs and Investigational Agents Identifies Patterns Associated with Gene and microRNA Expression. Mol Cancer Ther 2015;14:2452–62

36. Kaur G, Evans DM, Teicher BA, Coussens NP. Complex Tumor Spheroids, a Tissue-Mimicking Tumor Model, for Drug Discovery and Precision Medicine. SLAS Discov 2021;26:1298–314

37. Dexheimer TS, Coussens NP, Silvers T, Wright J, Morris J, Doroshow JH, Teicher BA. Multicellular Complex Tumor Spheroid Response to DNA Repair Inhibitors in Combination with DNA-damaging Drugs. Cancer Res Commun 2023;3:1648–61

38. Bliss CI. The Toxicity of Poisons Applied Jointly. Annals of Applied Biology 1939;26:585–615

39. Morris J, Kunkel MW, White SL, Wishka DG, Lopez OD, Bowles L, et al. Targeted Investigational Oncology Agents in the NCI-60: A Phenotypic Systems-based Resource. Mol Cancer Ther 2023;22:1270–9

40. Paull KD, Shoemaker RH, Hodes L, Monks A, Scudiero DA, Rubinstein L, et al. Display and analysis of patterns of differential activity of drugs against human tumor cell lines: development of mean graph and COMPARE algorithm. J Natl Cancer Inst 1989;81:1088–92

41. Pan R, Ruvolo V, Mu H, Leverson JD, Nichols G, Reed JC, et al. Synthetic Lethality of Combined Bcl-2 Inhibition and p53 Activation in AML: Mechanisms and Superior Antileukemic Efficacy. Cancer Cell 2017;32:748–60 e6

42. Hohtari H, Kankainen M, Adnan-Awad S, Yadav B, Potdar S, Ianevski A, et al. Targeting Apoptosis Pathways With BCL2 and MDM2 Inhibitors in Adult B-cell Acute Lymphoblastic Leukemia. Hemasphere 2022;6:e701

43. Decaudin D, Frisch Dit Leitz E, Nemati F, Tarin M, Naguez A, Zerara M, et al. Preclinical evaluation of drug combinations identifies co-inhibition of Bcl-2/XL/W and MDM2 as a potential therapy in uveal melanoma. Eur J Cancer 2020;126:93–103

44. Van Goethem A, Yigit N, Moreno-Smith M, Vasudevan SA, Barbieri E, Speleman F, et al. Dual targeting of MDM2 and BCL2 as a therapeutic strategy in neuroblastoma. Oncotarget 2017;8:57047–57

45. Wang GF, Liang E, Ming P, Rui L, Tang CY, Lv J, et al. Antitumor Activity of Dual BCL-2/BCL-Xl Inhibitor Pelcitoclax (APG-1252) in Natural Killer/T-Cell Lymphoma (NK/TCL). Blood 2021;138

46. Polley E, Kunkel M, Evans D, Silvers T, Delosh R, Laudeman J, et al. Small Cell Lung Cancer Screen of Oncology Drugs, Investigational Agents, and Gene and microRNA Expression. J Natl Cancer Inst 2016;108

47. Fairchild CK, Jr., Floros KV, Jacob S, Coon CM, Puchalapalli M, Hu B, et al. Unmasking BCL-2 Addiction in Synovial Sarcoma by Overcoming Low NOXA. Cancers (Basel) 2021;13

48. Isfort I, Berthold R, Heinst L, Wardelmann E, Larsson O, Trautmann M, Hartmann W. Interdependence of SS18-SSX-driven YAP1 and beta-Catenin Activation in Synovial Sarcoma. Mol Cancer Res 2023;21:535–47

49. Nakayama R, Zhang YX, Czaplinski JT, Anatone AJ, Sicinska ET, Fletcher JA, et al. Preclinical activity of selinexor, an inhibitor of XPO1, in sarcoma. Oncotarget 2016;7:16581–92

50. Liu Y, Azizian NG, Dou Y, Pham LV, Li Y. Simultaneous targeting of XPO1 and BCL2 as an effective treatment strategy for double-hit lymphoma. J Hematol Oncol 2019;12:119

51. Shang E, Zhang Y, Shu C, Ishida CT, Bianchetti E, Westhoff MA, et al. Dual Inhibition of Bcl-2/Bcl-xL and XPO1 is synthetically lethal in glioblastoma model systems. Sci Rep 2018;8:15383

52. Zhu ZC, Liu JW, Yang C, Zhao M, Xiong ZQ. XPO1 inhibitor KPT-330 synergizes with Bcl-xL inhibitor to induce cancer cell apoptosis by perturbing rRNA processing and Mcl-1 protein synthesis. Cell Death Dis 2019;10:395

53. Heisey DAR, Lochmann TL, Floros KV, Coon CM, Powell KM, Jacob S, et al. The Ewing Family of Tumors Relies on BCL-2 and BCL-X(L) to Escape PARP Inhibitor Toxicity. Clin Cancer Res 2019;25:1664–75

54. Zhang X, Wen XZ, Chen GJ, Zeng S, Men LC, Wang HB, et al. Phase I study results of APG-115, a MDM2-p53 antagonist in Chinese patients with advanced liposarcoma and other solid tumors. J Clin Oncol 2020;38

55. Qin A, Kalemkerian GP, Mohindra NA, Patel JD, Karapetis CS, Carlisle JW, et al. First-in-human study of pelcitoclax (APG-1252) in combination with paclitaxel in patients (pts) with relapsed/refractory small-cell lung cancer (R/R SCLC). J Clin Oncol 2022;40

