## Supplemental Material for "Drug combinations with apoptosis pathway targeted agents alrizomadlin, pelcitoclax, and dasminapant in multi-cell type tumor spheroids"

**Supplemental Table S1.** Multi-cell type tumor spheroid inoculation densities per well in 384-well microplates.

| Malignant cell line | Malignant cells per well | HUVEC[a] per well | hMSC[b] per well |
|---|---|---|---|
| 327498-153-R-J2 | 625 | 260 | 156 |
| 993429-296-R-J1 | 625 | 260 | 156 |
| SYO-1 | 313 | 130 | 78 |
| HS-SY-2 | 313 | 130 | 78 |
| 755483-174-R-J1 | 313 | 130 | 78 |
| 772245-204-R-J1 | 1250 | 521 | 313 |
| Rh18 | 2500 | 1042 | 625 |
| 521955-158-R2-J5 | 625 | 260 | 156 |
| 521955-158-R6-J3 | 1250 | 521 | 313 |
| PANC-1 | 625 | 260 | 156 |
| 323965-272-R-J2 | 1250 | 521 | 313 |
| 556581-035-R-J1 | 2500 | 1042 | 625 |
| OVCAR-5 | 1250 | 521 | 313 |
| OVCAR-8 | 313 | 130 | 78 |
| 126254-015-R-J1 | 625 | 260 | 156 |
| 494315-158-R-J1 | 1250 | 521 | 313 |
| 317291-083-R-J1 | 313 | 130 | 78 |
| 596521-263-R-J1 | 1250 | 521 | 313 |
| MPNST | 5000 | 2083 | 1250 |
| 186277-243-T-J2 | 1250 | 521 | 313 |
| 439559-082-T-J2 | 2500 | 1042 | 625 |
| 463943-066-T-J2 | 2500 | 1042 | 625 |
| 519858-162-T-J1 | 2500 | 1042 | 625 |
| 616215-338-R-J1 | 625 | 260 | 156 |
| 171881-019-R-J1 | 2500 | 1042 | 625 |
| MCF7 | 313 | 130 | 78 |
| MDA-MB-231 | 625 | 260 | 156 |
| 885512-296-R-J2 | 313 | 130 | 78 |
| NCI-H1876 | 2500 | 1042 | 625 |
| NCI-H196 | 2500 | 1042 | 625 |
| NCI-H510A | 5000 | 2083 | 1250 |
| NCI-H1417 | 5000 | 2083 | 1250 |

[a]human umbilical vein endothelial cells
[b]human mesenchymal stem cells

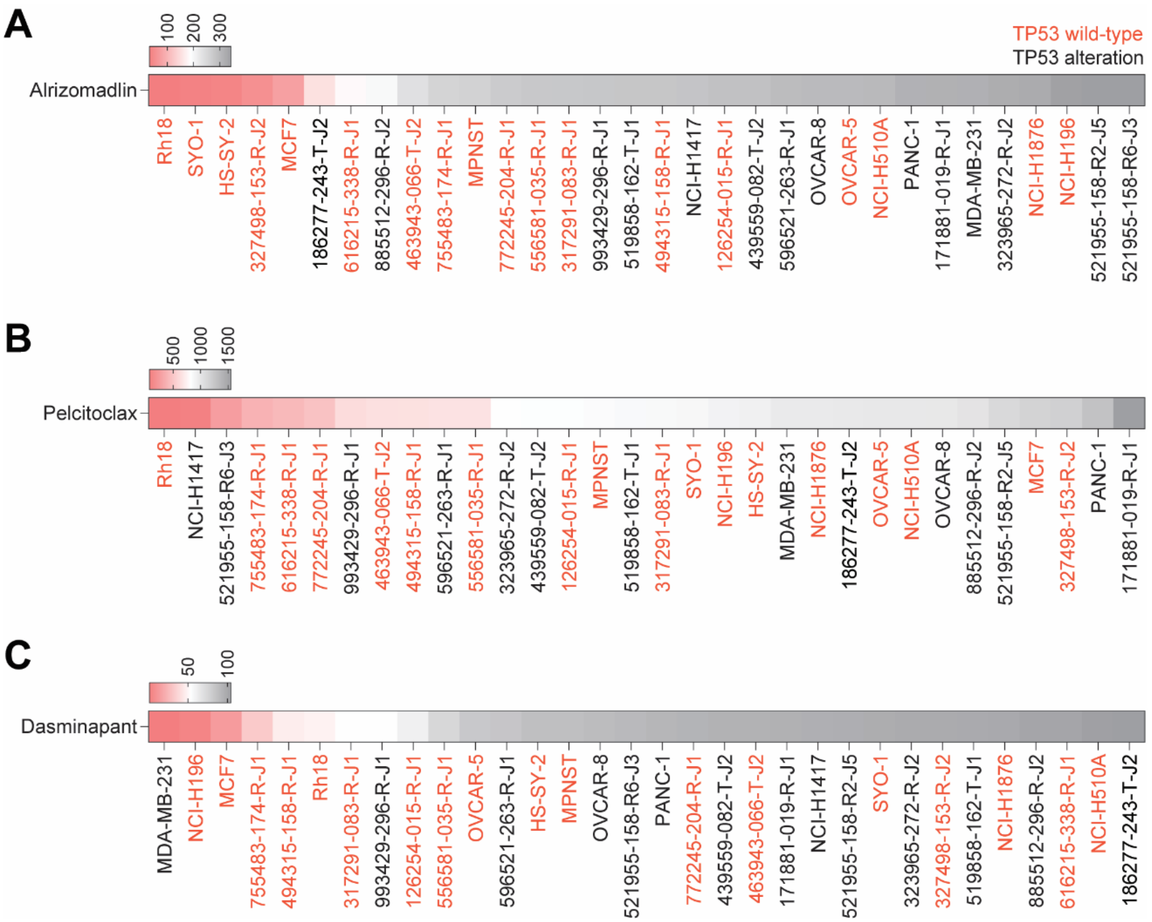


**Figure S1**. **Area under the concentration-response curve (AUC) for alrizomadlin, pelcitoclax, and dasminapant** (*see* **Figure 1**). Heatmaps depict the area under the curve (AUC) for A) alrizomadlin, B) pelcitoclax, and C) dasminapant, where red represents low AUC values and gray represents high AUC values. The text color of the cell line name indicates TP53 status, with red indicating wildtype and black indicating an alteration.

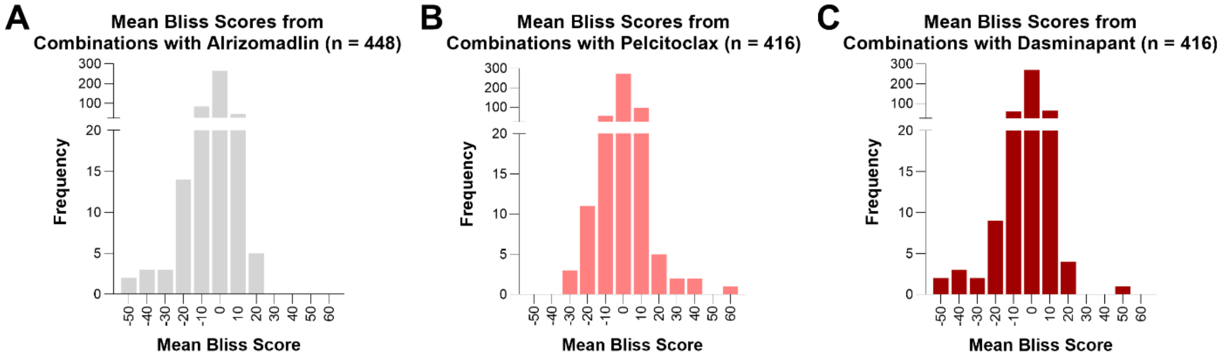


**Figure S2**. **Summary of mean Bliss independence matrix scores from all combinations of agents tested in thirty-two spheroid models**. A mean Bliss independence matrix score was calculated for each spheroid model from each combination's concentration matrix ([5 concentrations of drug A, $n \geq 3$ technical replicates] × [6 concentrations of drug B, $n \geq 3$ technical replicates] = [30 combination concentrations]) (*refer also to* **Table S3**). Histograms show the distribution of mean Bliss independence matrix scores for combinations with A) alrizomadlin ($n = 448$), B) pelcitoclax ($n = 416$), and C) dasminapant ($n = 416$).
